## Supplemental data for "Characterisation of *lmx1b* paralogues in zebrafish reveals divergent roles in skeletal, kidney, and muscle development"

***lmx1ba* and *lmx1bb* paralogues play tissue specific roles during zebrafish development**

**SUPPLEMENTAL TABLES AND FIGURES**

**Supplemental Table 1**

| | **Total fish** | **Otolith no.** | | | **Normal?** | | **Normal as a %** | |
|---|---|---|---|---|---|---|---|---|
| | | **2** | **1** | **0** | **Y** | **N** | **Y** | **N** |
| ***wt*** | 19 | 19 | 0 | 0 | 19 | 0 | 100% | 0% |
| ***lmx1ba***$^{-/-}$ | 28 | 28 | 0 | 0 | 27 | 1 | 96% | 4% |
| ***lmx1bb***$^{-/-}$ | 17 | 17 | 0 | 0 | 10 | 7 | 59% | 41% |
| ***dKO*** | 23 | 23 | 0 | 0 | 11 | 12 | 48% | 52% |

**Supplemental Table 1 – Otolith number is unaffected, but morphology is altered in *lmx1b* mutants compared to *wt* at 5dpf.**

**Supplemental Figure 1**

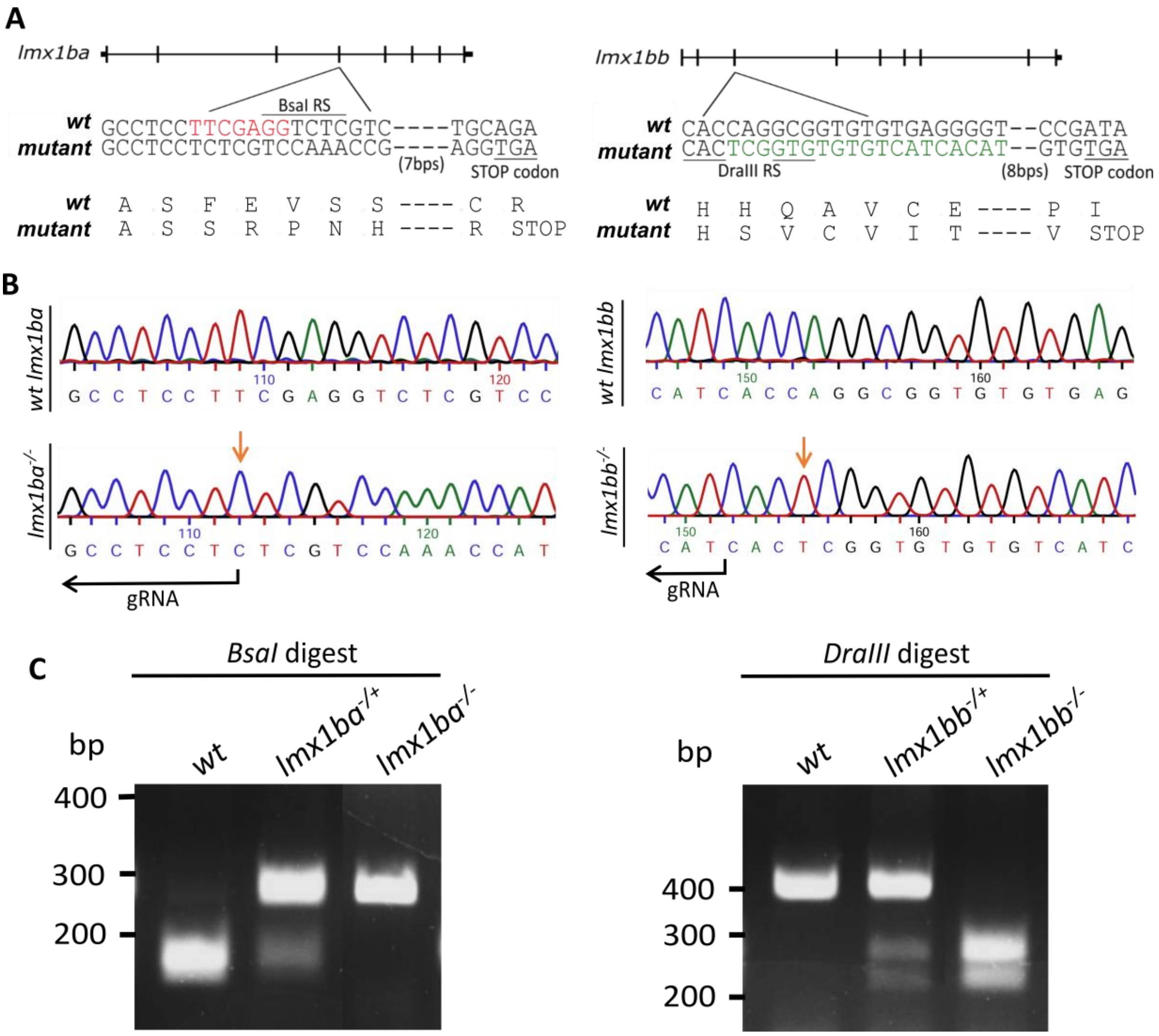


**Supplemental Figure 1 – *lmx1ba* and *lmx1bb* null fish lines generated using CRISPR-Cas9. (A)** Schematic showing exon position of selected mutations in both *lmx1ba* and *lmx1bb* populations, with resulting nucleotide deletion or insertion depicted in red and green, respectively, and subsequent amino acid changes leading to a stop codon. These nucleotide changes result in loss of the *BsaI* restriction site in *lmxba*$^{-/-}$ and gain of the *DraIII* restriction site in *lmx1bb*$^{-/-}$. Location and sequence of restriction sites used to genotype each population highlighted. RS, restriction site. **(B)** Representative *wt* and homozygous DNA traces of *lmx1ba* and *lmx1bb* from Sanger sequencing of DNA from F2 population larvae. Beginning of mutated sequences highlighted by orange arrow. For *lmx1bb* sequences, primer set 1 was used. **(C)** Example gel images showing restriction digest of *lmx1ba* and *lmx1bb* DNA with *BsaI* and *DraIII* restriction enzymes, respectively, to separate the genotypes of each population. *DraIII* restriction digest only possible with *lmx1bb* primer set 1.

**Supplemental Figure 2**

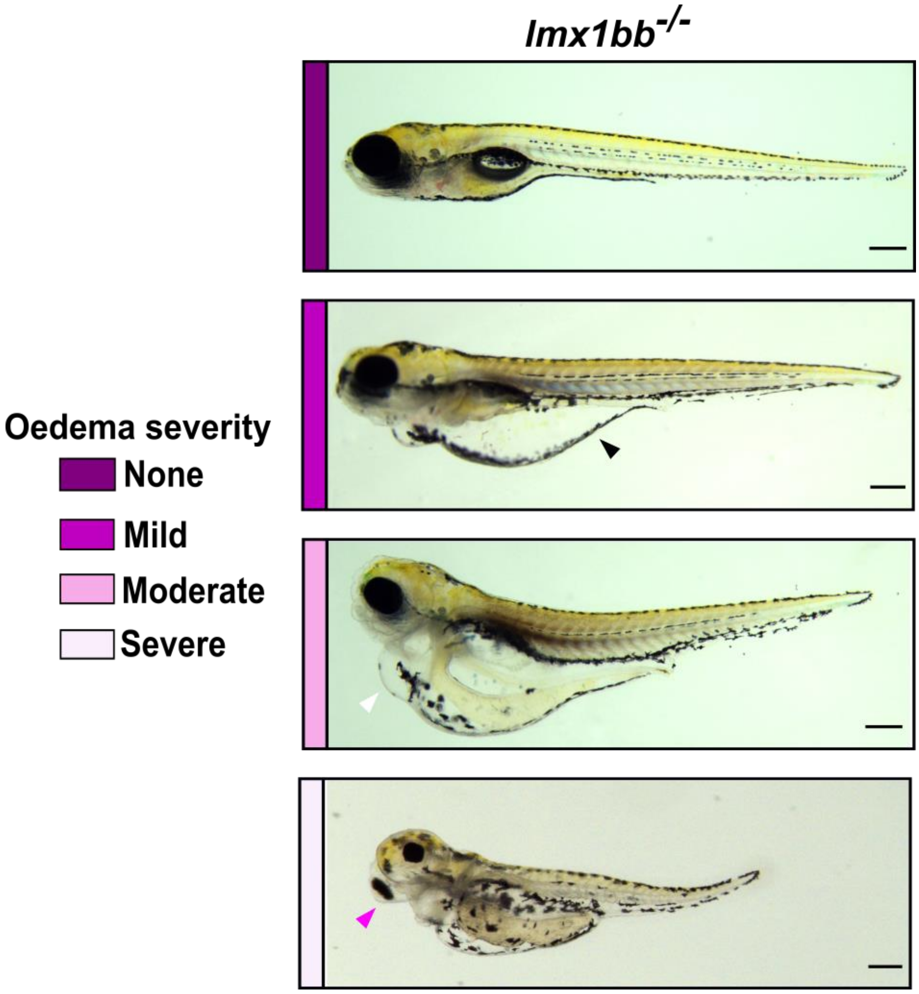


**Supplemental Figure 2 – *lmx1bb* mutants show range of oedema severity at 5dpf.** Stereomicroscope images of *lmx1bb* mutant larvae at 5dpf showing different severities of oedema. Mild oedema detected primarily as yolk sac oedema (*black arrowhead;* image same as one used in Figure 1B for *lmx1bb*$^{-/-}$); moderate oedema classified by onset of pericardial oedema (*white arrowhead*); severe oedema indicated by addition of eye oedema (*pink arrowhead*). Scale bar = 250µm.

**Supplemental Figure 3**

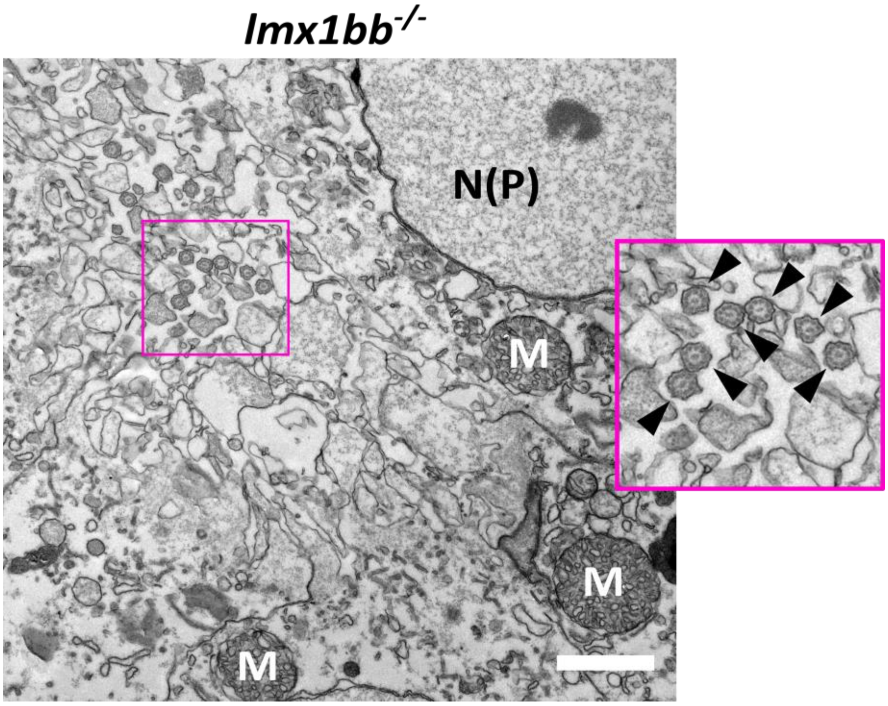


**Supplemental Figure 3 – Electron micrograph image of the glomeruli of a *lmx1bb* mutant at 6dpf.** In some areas, the *lmx1bb* mutants show complete loss of glomerular structure and organisation and swollen mitochondria. Inset shows accumulation of motile cilia (*black arrowheads*) in *lmx1bb* mutant which could be due to reduced kidney function. M = mitochondria; N(P) = nucleus of podocyte cell. Scale bar = 500nm.
